# Genotyping of paired clinical isolates using *PvCSP, PvMSP3*α*, PvMSP3*β and exploring STRs to differentiate between relapse and reinfection in *P. vivax*

**DOI:** 10.1101/2024.04.16.589389

**Authors:** Deepali Savargaonkar, Renuka Gahtori, Swati Sinha, Preeti Kumari, Paras Mahale, Bina Srivastava, Veena Pande, Himmat Singh Pawar, Anupkumar R Anvikar

## Abstract

The challenge of eliminating Vivax malaria is due to the relapses caused by hypnozoites. Despite several attempts to identify molecular markers to differentiate between relapse and new infection, a reliable marker has not yet been established. To address this issue, a genomic study was conducted on paired samples of patients who had experienced Plasmodium vivax infection twice. Genotyping was performed on paired samples from 10 vivax malaria patients using five molecular markers to distinguish between relapse and new infection. Our findings indicate that one sample represented a second episode that was a relapse of the first, while another sample had three episodes, two of which were relapse episodes. We combined our clinical records with molecular inferences to identify each pair as a relapse. This particular study provides a momentary view of forthcoming research endeavors that could be undertaken to distinguish between a relapse and a novel infection. Given the notable genetic variability of Plasmodium vivax, it is crucial to harness various markers to discern between relapse and new infection. The findings of this study are poised to be of immense utility for upcoming marker research in the aforesaid aspect.

## Introduction

In humans the disease is caused by five different species of Plasmodium which are *P. falciparum*, *P. vivax*, *P. malariae*, *P. ovale* and *P. knowlesi*. According to the world malaria report approximately 247 million malaria cases were reported from the 84 countries in year 2021 across the globe. (1). The number of malaria cases are significantly falling in the last two decades in South-East Asia Region i.e., reduced by 83.0% (1). In 2021, India reported a total of 0.16 million confirmed malaria cases and 90 deaths due to malaria (2). Among all malaria parasites, *Plasmodium falciparum* (*Pf*) and *Plasmodium vivax* (*Pv*) are the most commonly found species. In India, epidemiology of malaria is complex due to variable eco-epidemiological profiles, and different transmission factors. India remains a major contributor of *Pv* cases (83%) globally (3).

In recent years, there has been a surge in attempts to regulate malaria. These circumstances have resulted in a contradictory escalation in the proportion of P. vivax infections in diverse regions that have conventionally been recognized for their elevated levels of endemicity, such as Sri Lanka, Thailand, and Brazil. The fundamental reason behind this pattern can potentially be attributed to the fact that previous measures focused on the prevention and control of the infection, without duly considering the distinctive characteristics of P. vivax, which might have unintentionally contributed to the accumulation of sources of P. vivax infection (4).

*P. vivax* persist as dormant stage “hyponozoites” in liver cells and may cause disease after weeks to months of infection called relapse. Relapse study is challenging due to the lack of establishment of in-vitro cultures, and limited availability of functional assays.(5) Lack of reliable marker to distinguish relapse from reinfection complicates the aforesaid problems additionaly ()(6). Several molecular markers have been used for this purpose which primeraly includes P. *vivax Circumsporozoite Surface Protein* (*Pv*CSP)*, P. vivax Merozoite Surfavce Protein 3 alpha* (*Pv*MSP 3α)*, P. vivax Merozoite Surface Protein 3 beta* (*Pv*MSP 3β)*, P vivax Apical Membrane Antigen* etc (*Pv*APA)). (7,8,9)

The circumsporozoite protein (CSP) being the major cell surface protein of the *Plasmodium* sporozoites plays several roles such as maturation of sporozoites, invasion of salivary gland and hepatocyte inside mosquitoes and human host respectively (,10. The *P. vivax* CSP (*Pv*CSP) has three distinct domains which includes highly conserved domain at both the ends and a central domain with varying numbers of tendem repeats. Based on variation of sequences in the central domain, there are three types of the variant; the VK210 and VK247 and *P*. *vivax* like (11). The CSP gene has been targeted as the marker for differentiation of relapse from re-infection because of its highly polymorphic nature.

The erythrocytic stage starts with the invasion of *Plasmodium* in the RBC, this invasion is facilitated by a group of proteins one of such proteins in *Pv* is Merozoites Surface Protein (*Pv*MSP) which is encoded by several genes such as *Pv*MSP-1, *Pv*MSP3α, *Pv*MSP3β, *Pv*MSP3γ, *Pv*MSP4, and *Pv*MSP5 is encoded by genes *viz. Pv*MSP-1, *Pv*MSP3α, *Pv*MSP3β, *Pv*MSP3γ, *Pv*MSP4, and *Pv*MSP5 (13).

Polymorphism in *Pv*MSP3α and *Pv*MSP3β genes have been widely suggested for epidemiological studies as well as also used as a marker to differentiate between relapse and re-infection in the *P.vivax* (14). *Pv*MSP3α is also a potential vaccine candidate because of its association with immunogenicity (15).

The lack of reliably classifying the patient outcomes, specifically to differentiate between recrudescence, from reinfection or relapse poses challenge in clinical assessment of drug efficacy in *Pv* malaria (16).

In the present study, we have performed the data mining of *Pv* reference strain (i.e., Salvador I) genome followed by screening of minisatellite markers. Therefore, besides *PvCSP, PvMSP3*α and *PvMSP3*β, two newly screened minisatellite markers i.e., Ch05 and Ch14 have been used for differentiating relapse from re-infection in the paired samples from *Pv* infected patients.

## Material and Methods

### 1.1 Ethical approval

Ethical approval from Institutional Ethics Committee (IEC) was taken prior to collection of the samples for the study. Study was carried out using protocol which was reviewed and approved by the IEC (project Id-ECR/NIMR/EC/2017/90).

### 1.2 Patient record and Treatment

Fever Clinic of ICMR-National Institute of Malaria Research (ICMR-NIMR) perform routine diagnosis of malaria suspected individuals using microscopy. For all the confirmed vivax malaria patients, history of fever, age, gender and past malaria history were recorded. Malaria patients were treated according to the national drug policy (). A qualitative Glucose-6-Phosphate Dehydrogenase (G6PD) deficiency test was performed before administering primaquine. Treatment for vivax infection was given as per National treatment guideline (17). Vivax malaria patients with recurrent malaria episodes were identified using clinical Id of patient, name of patient, recorded previous history of malaria. Demographic details of vivax malaria patients, diagnosis of malaria, antimalaria treatment given were recorded using Microsoft excel 2007.

The genomic study was done on the samples collected from nine malaria patients during two vivax malaria episodes and in one patient during three malaria episodes. The aim of the study is to identify cases of relapse or new infection in paired samples (n=10), using molecular markers as well as on the basis of recorded details of primaquine intake and transmission or non-transmission season during the recurrent episodes. The months, December to June have been reported as months of non-transmission while July to November as transmission months for vivax malaria (,18).

### 1.3 Dried Blood Spots preparation

The blood samples from confirmed *Pv* malaria patients were collected by finger prick method aseptically at the fever clinic of the ICMR-NIMR, New Delhi. Three spots of blood were collected on a 3 mm filter paper (Whatman International Ltd., Maidstone, UK) for DNA isolation.

### 1.4 DNA isolation and diagnostic PCR for species confirmation

The genomic DNA was extracted from the blood samples collected on filter paper using the QIAamp DNA Blood Mini Kit (Qiagen, Hilden, Germany) as per the manufacturer’s instruction. To confirm the mono infection of *P. vivax* a nested PCR using genus-specific primers-specific for 18s rRNA as described by Johnston *et al* was performed (,19).

### 1.5 Genotyping of paired samples using *Pvcsp* gene

A nested PCR was performed in all the *Pv* positive samples to amplify the CSP gene using the sets of primers as described by Imwong *et. al*.(,20 (Table 1).

#### Analysis of the amplification product

The successfully amplified nested products (700bp) were further digested with *AluI* and *BstNI* enzymes and run on 2% agarose gel to detect the type of variants on the basis of banding pattern.

#### Genotyping of paired samples using Pvmsp 3**α** gene and Pvmsp 3**β** gene using RFLP enzymes

PvMSP 3α and Pvmsp 3β gene marker genes were amplified in all the paired samples by using the previously published protocol with some minor modifications in thermal profile. (8,21,22). Amplified nested PCR product was further digested with AluI, HhaI for msp3α and *PstI* for msp3 beta respectively.

Short tandem repeats (STRs) are highly informative genetic markers that have been used extensively in population genetics analysis to explore the genetic constitution of any gene or whole genome. They are an important source to explore the genetic diversity of any organism and can also have functional impact. For STR discovery in *P. vivax* Tandem Repeats Finder (TRF) was used with the default settings. STRs were filtered to remove mono-nucleotide repeats, percentage (%) match less than 90, % indels greater than 2, copy number less than 10 and greater than 20. Based on the period size, STRs were divided into micro (period size 2-7) and mini satellites (period size 8-20). Primer3 program was used to design primers for the STRs. The workflow for the analysis is as follows.

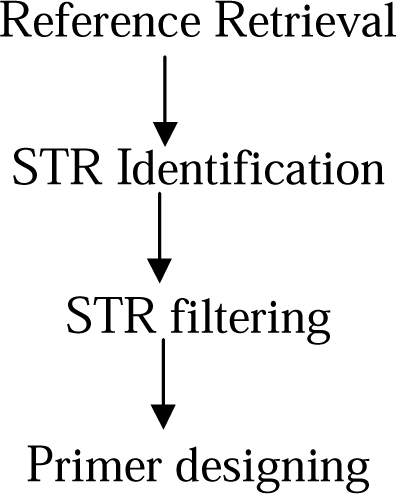

## Results

A total of 10 patients of *P.vivax* malaria with recurrent infection were included in the study. Seven out of these were male and 3 were females. The age ranged between 2.4 years to 52 years. The duration between two episodes of vivax infection was as low as six months to 1 year 22 days

### Data mining using bioinformatic tools in finding new molecular markers; Micro and minisatellite markers by exploring genome of *P. vivax-*

A total of 29,131 minisatellites and microsatellites were identified using TRF version 2. Among which 26,920 were found to be mono-nucleotide repeats during STR filtering, on applying filters period size in between 31%-89%, percent indels <3, copy number (10–20), minisatellites (period size 8-30), Microsatellites (period size 2-7). W found 469 and 137 mini and microsatellites that passed the aforesaid criteria. Finally, after primer designing, we were left with 264 and 101 mini and microsatellites respectively. Among all 14 chromosomes, chromosome no.14 had the highest no. of STRs reaching up to 3500 while the lowest was observed in chromosome no. 2 up to 2000 STRs (fig. 1). After applying filters, chromosome 12 and 14 still had the maximum no. of STRs i.e above 45 and 80 respectively. Whereas, chromosome no.1, 2, 3 and 4 had 20-40 no of STRs (fig 2).

**Figure 1:**
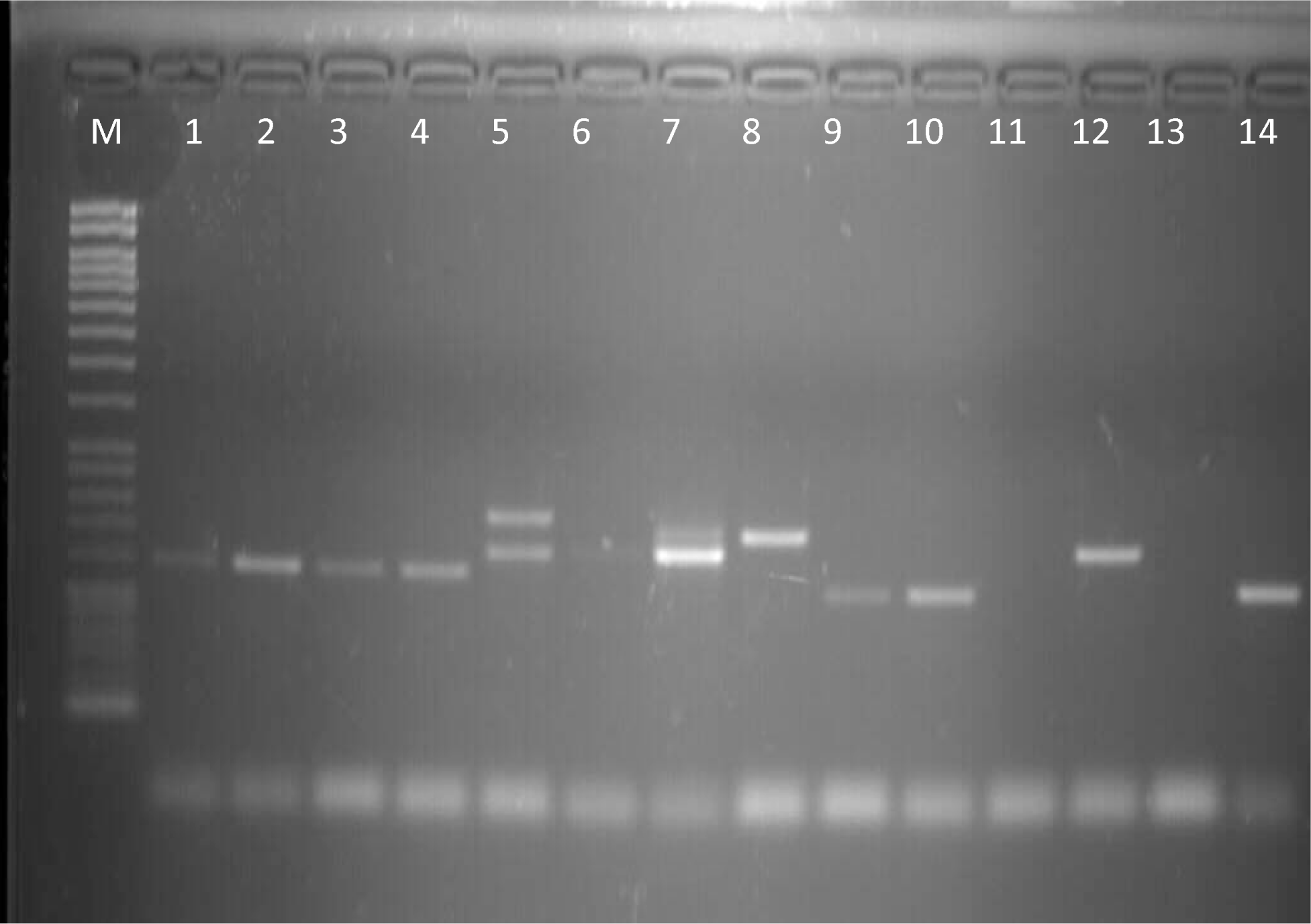
Agarose gel image showing amplification of Ch14 in the samples where M is DNA ladder of 50 base pairs and 1-14 are the samples of *Plasmodium vivax*

**Figure 2:**
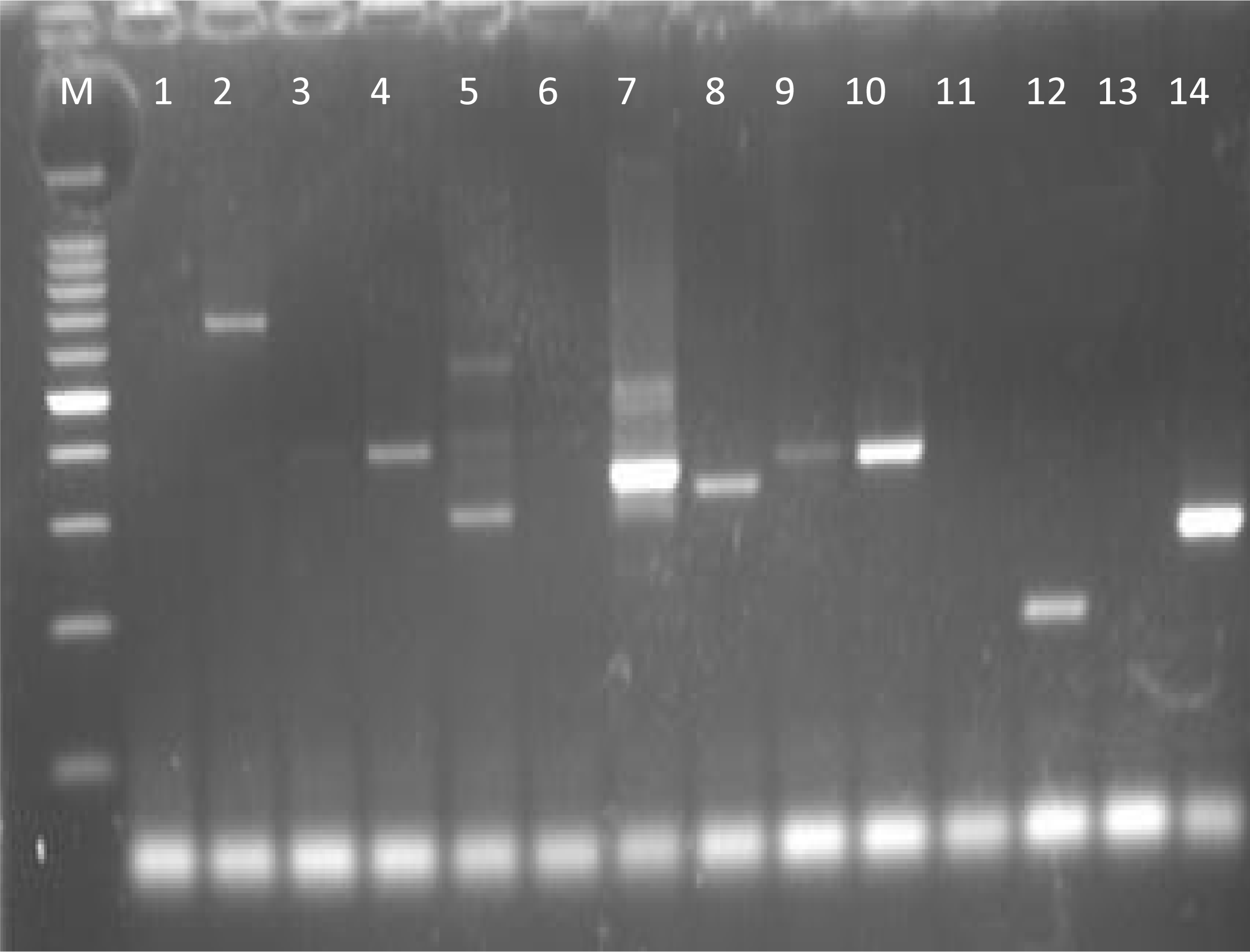
Agarose gel image showing amplification of Ch05 in the samples where, M is DNA ladder of 50 base pairs and 1-14 are the samples of *Plasmodium vivax*

For chromosome no. 14, approximately 50 STRs were found for which primers were designed. Chromosome no. 12 had approximately 45 STRs with primers (fig 3). The no. of STRs obtained during the analysis shows that after experimental validation there is a possibility to obtain markers to differentiate between relapse and reinfection.

**Figure 3:**
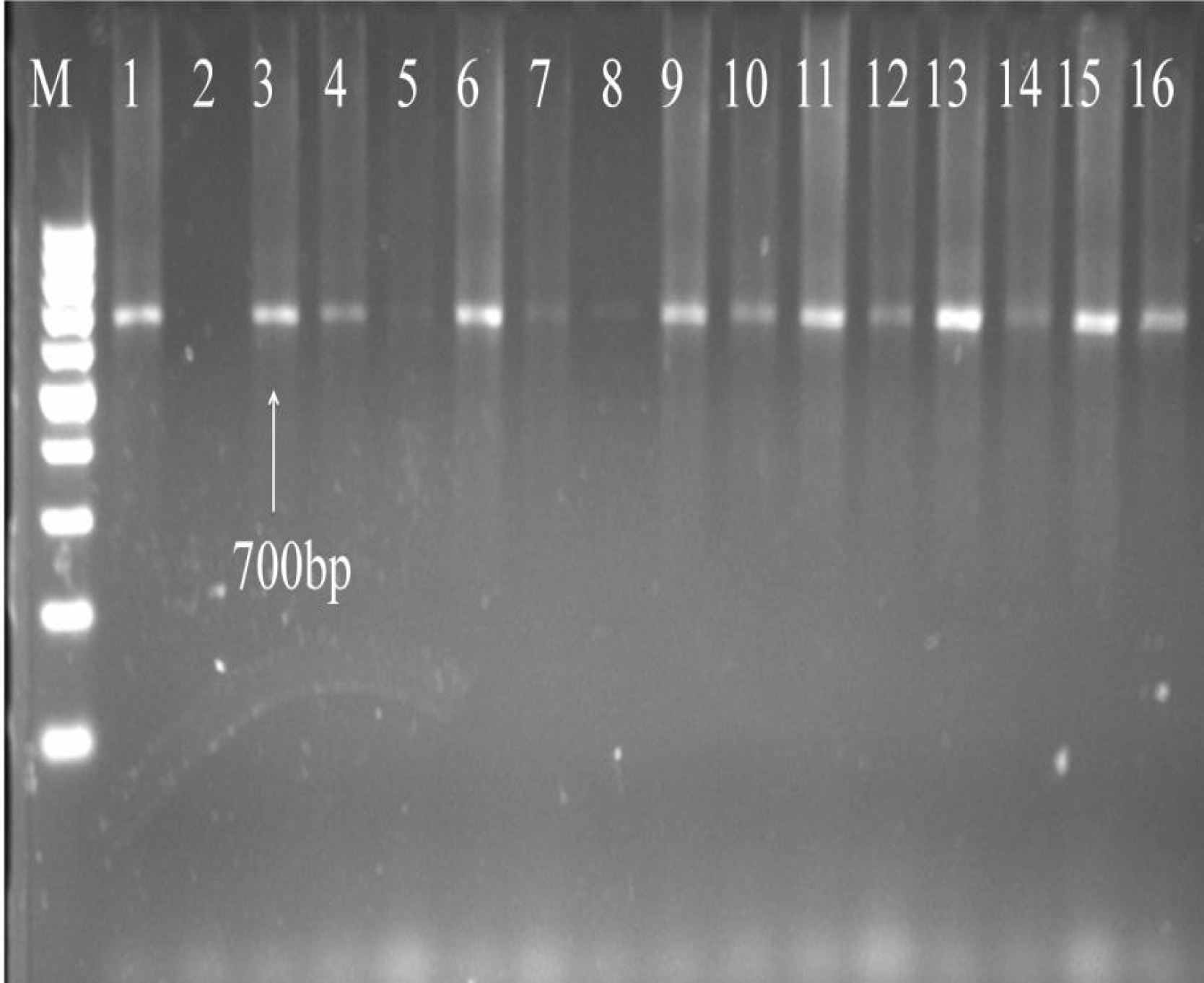
Agarose gel electrophoresis showing undigested nested product for *CSP* gene of *Plasmodium vivax* where M is DNA ladder of 100 base pair and 1 to 16 are samples.

Two minisatellites were applied to the samples named on the chromosome locations Ch 05 and Ch14. The Ch 05 was more polymorphic and resulted in a band size that varied from 220 to 650 base pairs whereas Ch14 had less variation in the band size (fig. 4 and 501).

**Figure 4:**
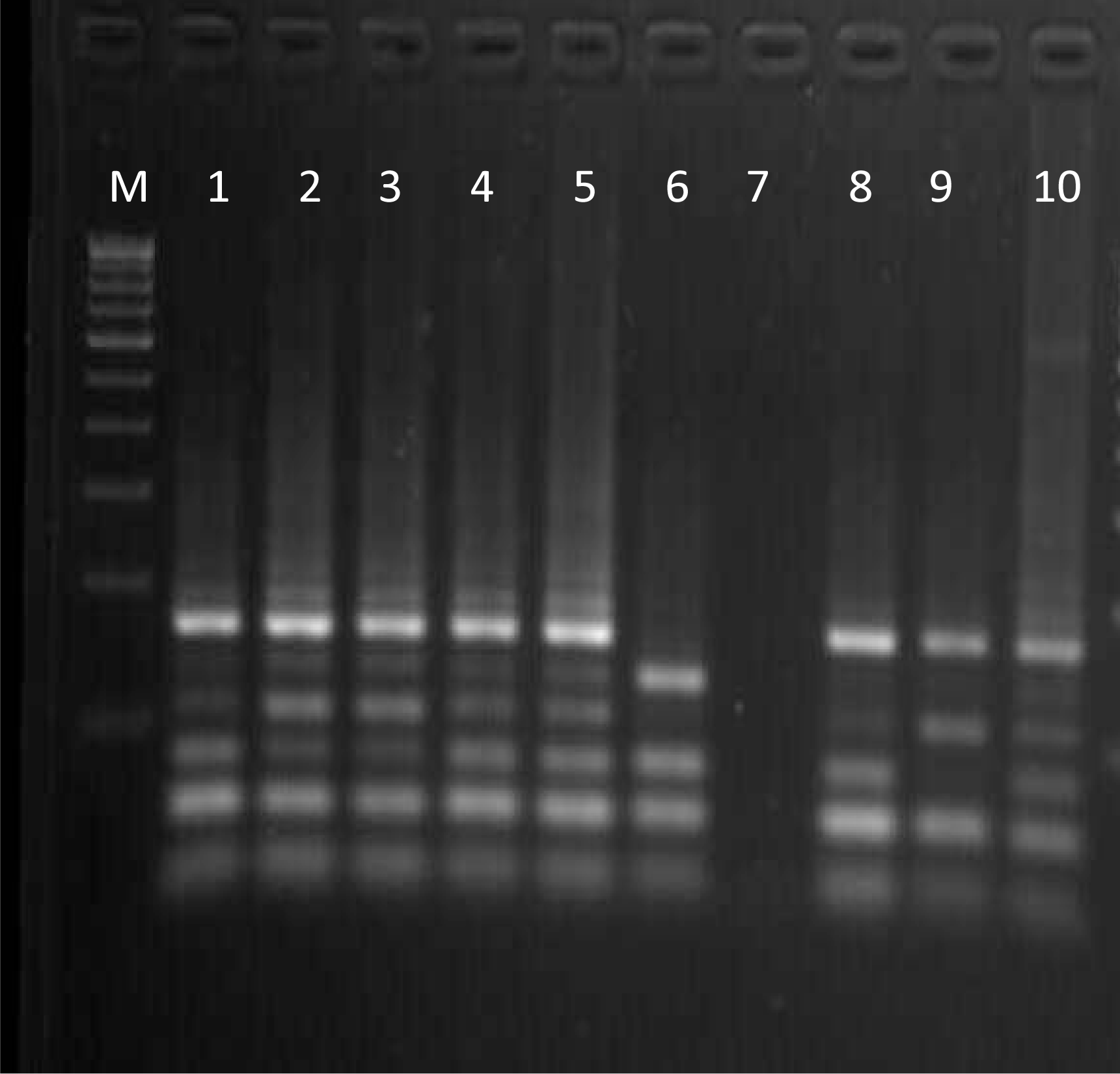
Agarose gel electrophoresis showing digested nested product for *CSP* gene of *Plasmodium vivax* where M is DNA ladder of 100bp and 1-10 are clinical samples.

### Genotyping

#### *Pv*CSP gene

All the paired samples were of VK210 type which is the most common genotype around the world. (Fig. 6 and 7). The sequences when N-blast on the NCBI database was confirmed to be VK210 genotype.

#### *Pv*MSP 3α gene and *Pv*MSP 3β gene

The nested product for *Pv*MSP 3α was 1.8 kb for nineteen samples whereas the other two samples showed band at 1.4 kb. When digested with *AluI* resulted in 12 different alleles (table 2) whereas*the HhaI* digestion resulted into 14 different allele types in the samples and both resulted in 26 different genotypes in these samples (table 3). The RFLP pattern of *P. vivax* isolates showed *AluI* is a better enzyme to uncover most of the variations compared with *HhaI* as stated by Prajapati et al (8).

The nested product for MSP 3β gene varied between 1.4 kb to 1.8 kb. Among all 1.8 kb was most frequent followed by 1.5 kb band. Only one sample showed up with 1.4 kb band (table 4).

### Clinical results

Interpretation of relapse or new infection was made by considering the results by molecular marker as well as recorded details of primaquine intake and transmission or non-transmission season during recurrent episode. Out of 11 paired samples, 6 samples were interpreted as confirmed new infection while 5 were interpreted as relapse out of 7 parameters, at least 4 parameters showed concordant results. Among these paired samples, for two paired samples, all 5 molecular markers gave concordant results while for another 3 paired samples, 4 molecular markers gave concordant results. We also documented the history of travel within 1 month. During transmission season, four patients with new infections and had a travel history of these cases could be imported cases. They had travelled to districts of Uttar Pradesh.

**Case 1**- All the markers showed same banding pattern. The period between two episodes was six months nine days and the second episode occurred in the non-transmission season. The patient had not taken primaquine during the primary attack. Since the clinical data was in concordance with the molecular marker analysis; it was concluded as a relapse case.

**Case 2**- Only one marker showed a difference in bands of MSP3β. The recurrent episode occurred in the non-transmission season and the period between the episodes was more than one year. It was considered as a relapse case.

**Case 10**- Four-year-old girl, had three episodes. She had not taken primaquine. The second episode was within 8 months of the primary attack occurred in non-transmission season whereas the third episode occurred 11 months after the primary attack and during transmission season. On the basis of the banding pattern of selected molecular markers, it was considered that the second episode was a relapse whereas the third was re-infection which might be caused because of the new mosquito bite.

In our study we have used two previously recommended markers i.e., *MSP* 3α and β and two newly developed minisatellites ch 05 and ch 14. Use of more than one marker could give a clearer picture. Out of ten patients, relapse occurred in two even after taking primaquine. We observed that all relapses occurred in non-transmission season i.e., from November to June (Table 5).

## Discussion

In India around 82 percent of population is susceptible to malaria. With the presence of various relapse phenotypes with varied latency durations, the epidemiology of *P. vivax* varies significantly across India and the hypnozoite reservoirs pose risk of re-transmission where it was supposed to be eradicated (23). Furthermore, accurately classifying patient outcomes poses a challenge, particularly when distinguishing between recrudescence, reinfection, or relapse in cases of *P. vivax* malaria recurrence during clinical evaluations and trials conducted to assess therapeutic effectiveness (15).

The genetic diversity of *P. vivax* is less explored because of lack of parasite culture methods which restrict to perform genetic studies on the organism and also due to the absence of more informative markers and is majorly based on few loci like Mt DNA and Duffy binding protein (,24).

Several studies have suggested different markers which are polymorphic and well suited for epidemiological studies as well as predicted to be probable marker to differentiate between relapse and new infection, henceforth, we studied previously established molecular markers like *CSP*, MSP *3*α and β (5, 8, 13,21,2,225,,26and also looked for minisatellite markers and selected two minisatellite markers which were newly developed as previously the use of minisatellite markers has shown promising results. Adding to it, two parameters primaquine intake and season (transmission or non-transmission at the time of the recurrent attack) were also considered in concluding the results. (5).

Out of 11 paired samples, only in two samples all the molecular markers and demographic criterias (primaquine intake and season) showed similar results. Due to the extensive genetic diversity in *P. vivax*, there is need to consider clinical, demographic parameters along with molecular markers to differentiate relapses from new infections. It was observed that during the transmission season, four patients with new infection had history of travel. They had travelled to districts of Uttar Pradesh (UP). These cases could be imported cases as according to the epidemiological situation, UP falls under category 2 while Delhi is under category 1 with <1 annual parasite incidence (16).

The duration between the primary attack and recurrent infection ranged from 6 month to 1 year and 22 days. As the duration was more than 30 days, the recurrent attacks could be either relapse or new infection and not recrudescence. We observed that all the relapses occurred in non-transmission season from November to June (17). In two patients, recurrence occurred even after intake of primaquine. This may be attributed to poor compliance to primaquine due to the long treatment course of 14 days (27).

In this study the purpose of using *CSP* marker was to screen the paired samples at first as if any of the paired samples would have different strains of *CSP* it would have either a mixed infection or a new infection to be appropriate. In the present study, VK210 genotype was predominant, which is the best adapted variant across the world (19) and is predominantly reported from South East Asia (Myanmar, 66%, Thailand, 77%), South Asia (Nepal, 95%), South America (Brazil, 86%), Central America (Honduras, 100%) countries. Infection with CSP variations can change the host’s medication responsiveness, illness intensity, and humoral response patterns (22,28). CSP polymorphisms influences the cytokine balance and the parasite burden in vivax malaria. Furthermore, the VK247 variation resulted in a more significant inflammatory profile as well as the highest parasite burdens, suggesting that these CSP polymorphisms have universal impacts on factors that may influence the course of infection (29).

Tandem repeats, also known as neutral genetic loci, are thought to be possible genetic markers for determining the population genetic makeup and evolutionary history of an organism. Numerous population studies have taken advantage of polymorphism in two categories of neutral loci: length polymorphism in micro- and mini-satellites and SNPs in so-called housekeeping genes. The *P. vivax* genome sequence has revealed a significant number of mini-satellites (31,30).

Genome-wide polymorphic microsatellite and minisatellite markers are frequently employed to identify the genetic diversity and population structure of human malaria parasites (31,32,33.

Using mini-satellite markers *the* genetic diversity of *P. vivax* on the Indian subcontinent revealed huge neutral genetic variation in Indian field isolates (35,34). Differentiating relapse from new infection will help in knowing the actual burden of vivax due to relapse and new infection. Reliable markers to differentiate between relapse and new infection will be useful to evaluate the efficacy of antimalarial drug. These PCR-RFLP subtyping methods on applying to massive epidemiological studies for molecular surveillance to understand genetic population of P. vivax and to supervise the genetic variation of the parasite circulating in the region as well as may be fruitful in standardizing a method to differentiate between relapse and reinfection.

## Conclusion

The use of a set of markers to differentiate between relapse and reinfection could give a much clearer picture in the endemic settings. The use of only one of them may create ambiguity. This study is a snapshot of the work to be done and highlights the need for well-documented patient records, follow ups and genotyping of paired samples on a large scale in order to know the true burden of vivax relapses. This approach will be useful in later stage of malaria elimination when during case-based intervention approaches.

## Acknowledgment

Authors are thankful to ICMR for providing Senior Research Fellowship (id 2019-5269) and Director ICMR-NIMR for the constant support.

## Contribution

DS and AA conceived the idea, DS managed the patients, obtained patient details and clinical samples, RG and SS designed the experiments, RG, SS and PK performed the experiments, RG and SS wrote the manuscript, RG and SS analyzed the data, PM, VP, HS, DS, BS, and AA reviewed the manuscript. All the authors read and approved the manuscript.

There is no conflict of interest among the authors.

## Legends for table and figures

**Table 1**: The list of primers used in study

**Table 2:** Msp 3 alpha gene and its digested product with *AluI*.

**Table 3:** Msp 3 alpha gene and its digested product with *HhaI*.

**Table 4:** Table for Msp 3 beta gene and its digested product with *Pst I*

**Table 5:** The consolidated results of the markers applied with the results obtained for different pateints

**Figure 1:** Number of STRs discovered per chromosome.

**Figure 2:** Number of STRs per chromosome after applying filters.

**Figure 3:** Number of STRs for which primers were found

**Figure 4:** Agarose gel image showing amplification of Ch14 in the samples where 1 is DNA ladder of 50 base pairs and 1-14 are the samples of *P. vivax*

**Figure 5:** Agarose gel image showing amplification of Ch05 in the samples where, 1 is DNA ladder of 50 base pairs and 2-15 are the samples of *P. vivax*

**Figure 6:** Agarose gel electrophoresis showing undigested nested product for *CSP* gene of *P. vivax* where M is DNA ladder of 100 base pair and 1 t0 16 are samples.

**Figure 7:** Agarose gel electrophoresis showing digested nested product for *CSP* gene of *P. vivax* where M is DNA ladder of 100bp and 1-10 are clinical samples.

